# γδ T cells are the prime anti-tumoral T cells in pediatric neuroblastoma

**DOI:** 10.1101/2025.01.23.634553

**Authors:** Suzanne M. Castenmiller, Anne L. Borst, Leyma Wardak, Jan J. Molenaar, Maria Papadopoulou, Ronald R. de Krijger, Alida F. W. van der Steeg, David Vermijlen, Rosa de Groot, Judith Wienke, Monika C. Wolkers

## Abstract

High-risk pediatric neuroblastoma patients have a dismal survival rate despite intensive treatment regimens. New treatment options are thus required. Even though HLA expression in neuroblastoma is low and immune cell infiltrates are limited, the presence of tumor infiltrating lymphocytes (TILs) is indicative for better patient survival. Here, we show that most tumor lesions contain viable immune cell infiltrates after induction chemotherapy, with high percentages of CD3^+^ T cells. We therefore expanded the TILs and tested their anti-tumoral activity. With sufficient starting material, TIL expansion was as efficient as for adult solid tumors. However, whereas TIL products from adult tumors almost exclusively contained αβ T cells, in neuroblastoma-derived TILs, γδ T cells expanded with similar efficacy as αβ T cells. Importantly, the anti-tumor responses in response to autologous tumor digest primarily originated from (Vδ1- and Vδ3-expressing) γδ T cells, and not from αβ T cells. In conclusion, this finding creates a window of opportunity for immunotherapy for neuroblastoma patients, with γδ T cells as potential prime responders.

## INTRODUCTION

Neuroblastoma is the most common extracranial solid cancer in children^1,2^. High-risk patients receive intensive treatment with debulking chemotherapy, surgical resection, and additional round(s) of high-dose chemotherapy followed by autologous stem cell therapy, radiotherapy, or the recently developed antibody-based anti-GD2 immunotherapy^3^. Yet, their 5-year survival rate does not exceed 50%^1,4^, highlighting the urgent need for novel treatment options.

Neuroblastoma is considered a ‘cold’ tumor with low mutational tumor burden, low human leukocyte antigen class I (HLA-I) expression, and low immune cell infiltrates^5–8^, which combined impedes the tumor cell recognition by T cells^9^. Nonetheless, the promising outcomes of anti-GD2 immunotherapy^10–12^ have sparked interest in utilizing the immune system to combat neuroblastoma^3,13^. Enhancing immune responses against neuroblastoma could indeed be powerful for improving the patients’ prognosis.

Adoptive therapy with tumor infiltrating lymphocytes (TIL therapy) is coming of age for treating solid tumors^14,15^. Reinfusion of in vitro expanded autologous T cells from tumor lesions achieved high response rates in a phase III clinical trial in melanoma patients with disseminated tumors^15^. Owing to this success and the recent FDA approval for TIL therapy for melanoma, TIL therapy is currently evaluated in other solid tumors, including non-small cell lung cancer^16–19^, renal cell carcinoma^20^, bladder cancer^21^ and ovarian cancer^22^. TIL therapy currently focusses on highly immune-infiltrated ‘hot’ tumors with a high mutational burden. Yet, clinical responses were also reported for TIL therapy in so-called ‘cold’ tumors with little to limited immune infiltrates and mutational burden, such as breast, ovarian, colorectal and pancreatic cancers^23–26^.

Even though neuroblastoma is considered a ‘cold’ tumor, high TIL density positively correlated with the patient outcome^7,27^. Chemotherapy, the standard-of-care for high-risk neuroblastoma patients promotes the influx of immune cells into tumors^28,29^. We therefore hypothesized that neuroblastoma lesions after induction chemotherapy contain tumor-reactive T cells that could potentially be utilized for therapeutic purposes. Here, we report effective generation of tumor-reactive TIL products from neuroblastoma tumor lesions. Importantly, the composition of pediatric TIL products substantially differed from those of TIL products from adult tumors. Indeed, γδ T cells were enriched in neuroblastoma lesions compared to adult tumors, and this enrichment was maintained throughout the expansion protocol. The γδ T cells expanded equally well as the αβ T cell receptor-expressing CD8^+^ or CD4^+^ T cells. Furthermore, γδ T cells were the prime T cell subset with anti-tumoral activity. Our findings thus uncover fundamental differences between adult and pediatric neuroblastoma derived anti-tumor responses.

## RESULTS AND DISCUSSION

### T cells constitute the majority of immune infiltrates in neuroblastoma lesions

We first studied the immune cell composition of neuroblastoma tumor lesions. We generated single cell suspensions from 20 tumor lesions obtained from 18 patients after debulking surgery (Table S1). On average, 32.7% of the cells were viable, of which a median of 28.3% ± 23.0% were CD45^+^ immune cells (Figure 1A-C; Figure S1A). Owing to the cryopreservation of tumor digests prior to analysis, myeloid cells such as monocytes and neutrophils were underrepresented (Figure S1A). Of the remaining immune cells, CD3^+^ T cells were with a median of 64.5% most abundant (Figure 1D). Monocytes, B cells, NKT cells, and NK cells were also detected, but with highly variable percentages ranging from 0% - 40.2% (Figure 1D; Figure S1A).

**Figure 1.**
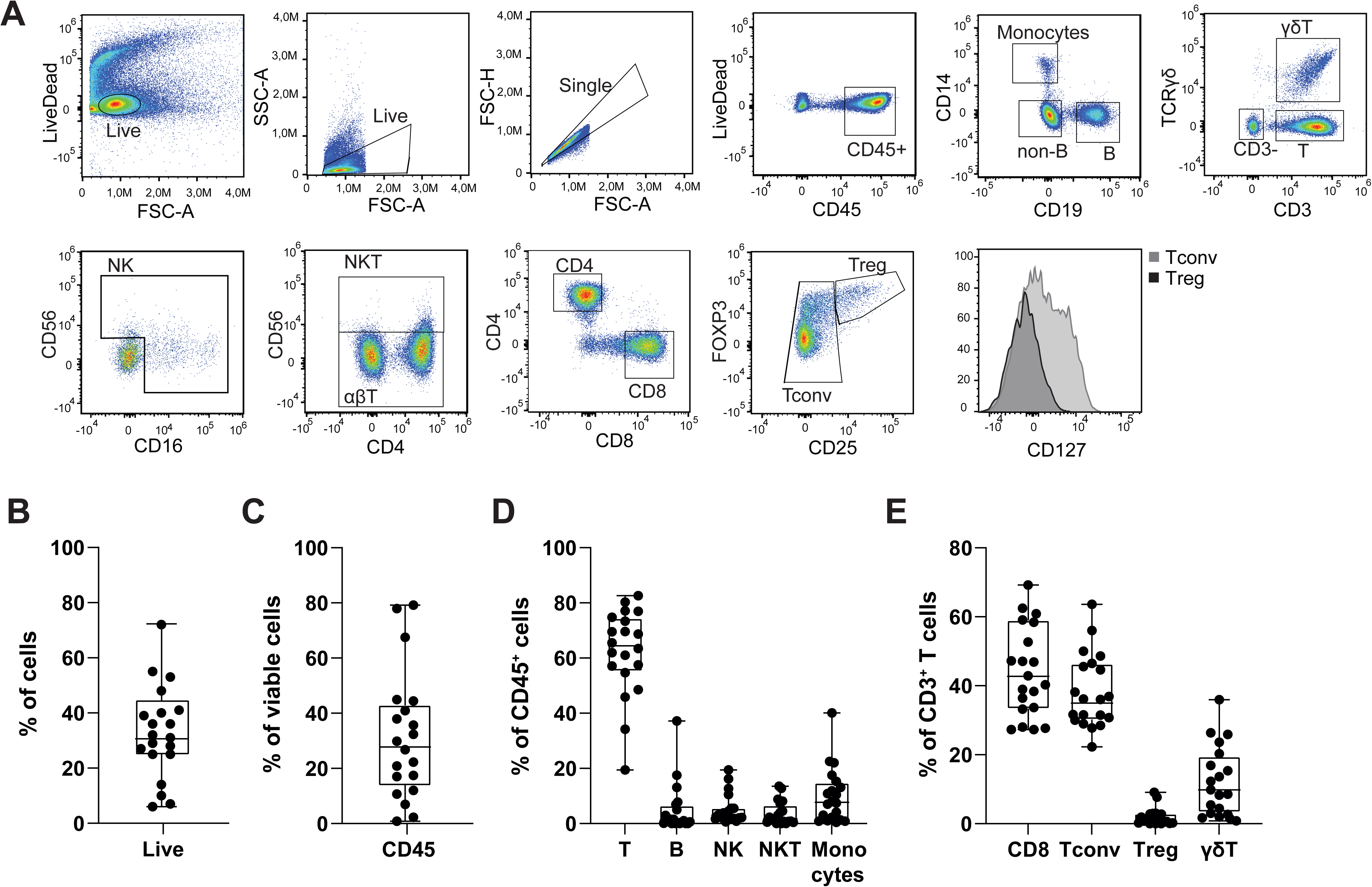
Immune cell composition of pediatric neuroblastoma tumor lesions. **(A)** Representative gating strategy for analyzing immune cell infiltrates (sample M189AAF_2). **(B-E)** Percentage of **(B)** viable cells in neuroblastoma tumor lesions, **(C)** CD45^+^ immune infiltrates of viable cells, **(D)** different immune cell subsets as percentage of total viable immune infiltrates, and **(E)** T cell subsets as percentage of total T cell infiltrate. n=20 samples, each dot represents one tumor sample. Box and whisker plots depict median, minimum and maximum values (whiskers) and 25th pct and 75th pct (box).

CD3^+^ T cell infiltrates in neuroblastoma lesions primarily consisted of αβ CD8^+^ T cells and conventional αβ CD4^+^ T cells (Tconv), together accounting for a median of 80% of CD3^+^ T cells (Figure 1E; Figure S1A). CD127^low^CD25^+^Foxp3^+^ regulatory T cells (Tregs) were detected in 17 out of 20 (85%) tumor lesions, yet with low percentages (Figure 1E; Figure S1A). We also detected γδ T cell infiltrates with a median of 9.2%, comprising up to 36% of the CD3^+^ T cells in one tumor lesion (Figure 1E; Figure S1A). Of note, the percentage of γδ T cells of CD3^+^ T cells reflect the percentages of γδ T cells reported in tissues of children^30^. The high variability in CD45^+^ cell infiltrates and its composition could not be attributed to different disease stages (LR; Low risk, MR; Medium risk, HR; High risk) or different treatment regimens (Table S1). In conclusion, even though pediatric neuroblastoma is considered a ‘cold’ tumor, almost all tumor lesions tested contain immune infiltrates, with high percentages of T cells.

### TILs can be effectively expanded from neuroblastoma lesions

Having established that T cells are present in neuroblastoma lesions, we tested their capacity to expand. A recent study reported successful TIL expansion from neuroblastoma lesions^31^, however limited tumor reactivity had been observed^31^. Since αCD3/αCD28 activation was used immediately for T cell expansion^31^, this protocol may have favored the expansion of T cells from contaminating blood, or of non-tumor reactive, bystander tissue resident T cells that are amply present in solid tumors^32,33^. This in turn may hamper the outgrowth of tumor-reactive, yet to some degree dysfunctional T cells^32,33^. We therefore used the Rapid Expansion Protocol (REP^34,35^), which for the first two weeks of culture (pre-REP) uses only the addition of recombinant human IL-2 for TIL expansion from tumor digests^15,36^. This culture setup allows for specific antigen presentation from the tumor cells in the pre-REP phase and is considered to support the survival and expansion of tumor-specific T cells^15,36^.

TILs from neuroblastoma lesions expanded on average 15-fold during the pre-REP phase, and 200-fold during the second two weeks with αCD3 and IL-2 (REP phase), resulting in a total expansion of ∼1500-fold (Figure 2A). However, the efficiency of TIL expansion was not uniform. Whereas 9 out of 20 (45%) TIL cultures expanded less than 500-fold (Figure 2A), 11 out of 20 (55%) TIL cultures expanded about 3000-fold (Figure 2A), which is comparable to the TIL expansion rate reported for adult tumors^15,18,19^. When we compared the patient’s age (in months) at the time of tumor resection, we found a slight association with the efficiency of TILs to expand (Figure 2B, left panel). Treatment intensity negatively correlated with TIL expansion (Figure 2B, middle panel). Most prominently, the number of cells that were available for starting the TIL cultures significantly correlated with TIL expansion (Figure 2B, right panel). Low cell numbers primarily stemmed from tumor needle biopsies as input material (Table S1). In sum, we conclude that sufficient starting material from tumor lesions is pivotal to achieve efficient TIL expansion.

**Figure 2.**
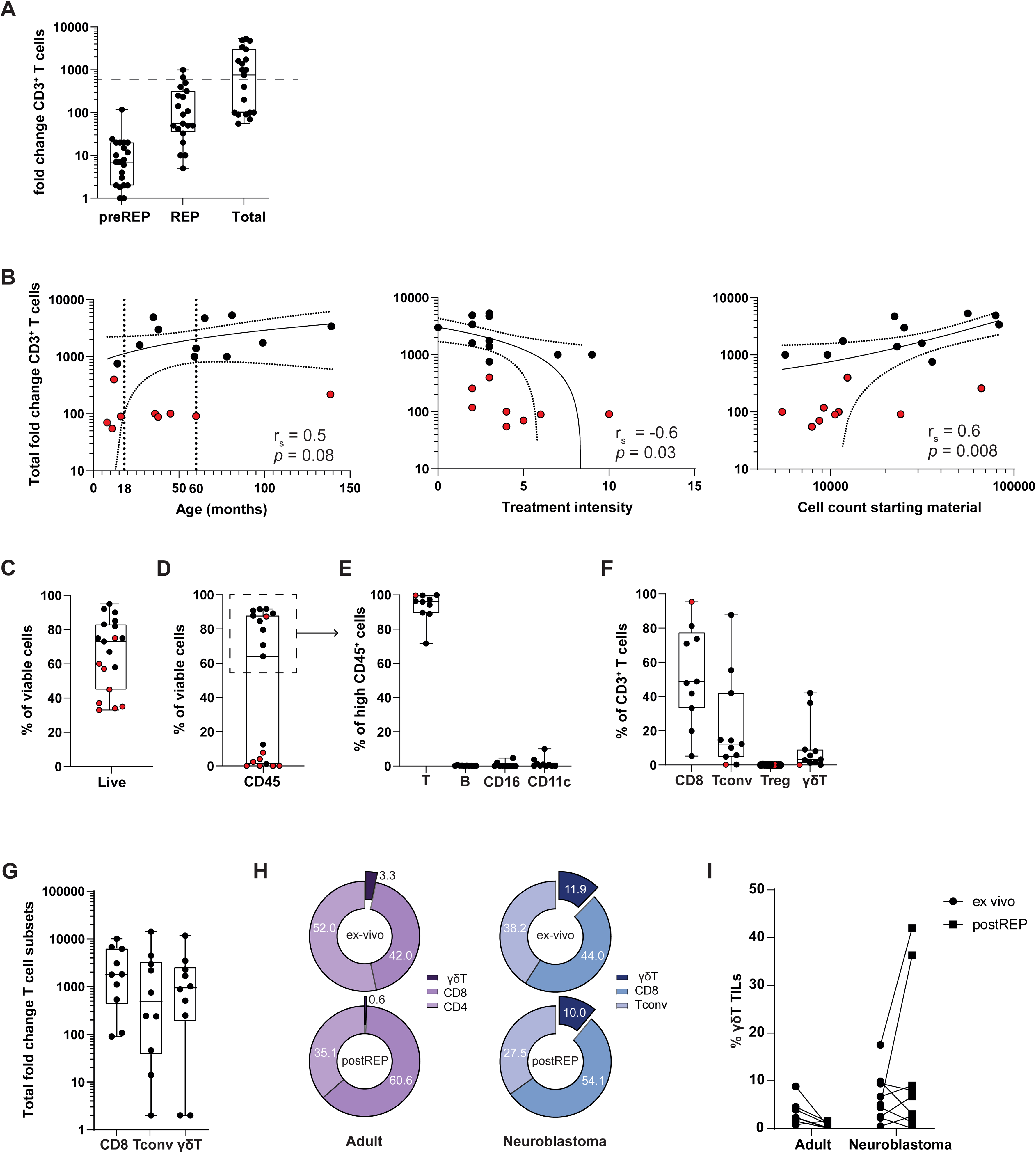
Expansion potential and composition of TIL products from neuroblastoma lesions. **(A)** Fold expansion of TILs during the first 10-13 days of the pre-Rapid expansion phase (pre-REP left), during the second 10-13 days (REP; middle) and pre-REP and REP combined (Total; right). Dotted line indicates 500-fold expansion, n=20. **(B)** Correlation plots between total expansion (y-axis) and patients’ age in month at time of tissue collection (left), treatment intensity based on total number of treatments received prior to surgery (middle), and total cell count of the starting material for TIL expansion (right). r_s_ and adjusted p-values are indicated in each plot. P values were corrected for six multiple comparisons using Bonferroni method. Red dots indicate expansion below 500-fold (n=9). **(C)** Percentage of viable cells in TIL products (n=20). **(D)** Percentage of CD45^+^ cells in TIL products (n=19, one sample was excluded due to insufficient expansion). **(E)** Percentage of indicated immune cell subsets as percentage of CD45^+^ cells and **(F)** T cell subsets as percentage of CD3^+^ cells. **(G)** Total fold expansion for CD8^+^, Tconv and γδ T cell subsets. **E-G:** n=10; only TIL products with high percentages of CD45^+^ cells were included **(H)** Indicated T cell subsets as percentage of CD3^+^ T cells ex-vivo (n=20) and after REP (n=10). **(I)** γδ T cell content as percentage of CD3^+^ T cells ex-vivo (dots) and upon REP (squares) for adult tumor samples (melanoma n=2, non-small cell lung cancer n=3, renal cell carcinoma n=3) and for neuroblastoma (n=10). Each dot represents one sample. Box and whisker plots depict median, minimum and maximum values (whiskers) and 25th pct and 75th pct (box). Correlation plots depict the simple linear regression line with the 95th pct coincidence interval (dotted line).

We next phenotyped the TIL products after REP culture (Figure S1B, S1C). Cell viability of the 19 measured TIL products was on average 65.84%, yet with high variability. Well-expanding TIL products (>500-fold) reached a mean of 82.5% viability on average, which is above the threshold for clinical use (Figure 2C, black dots; Figure S1B). Poorly expanded TIL products (<500-fold) contained low numbers of CD45^+^ cells (0–12.5%). In contrast, well-expanding TIL products contained 64.0–91.7% CD45^+^ cells, and in 9 samples >90% of the CD45^+^ cells were CD3^+^ T cells (Figure 2D, 2E, black dots; Figure S1B). Only two TIL products with <100-fold expansion contained high percentages of viable cells (Figure S1D, *colored dots*), of which one contained almost exclusively CD45^+^CD3^+^CD8^+^ T cells (Figure S1D, *blue dot*). CD19^+^ B cells and Foxp3^+^CD4^+^ T cells were absent in well-expanding TIL products, and only 2 out of 10 well-expanding TIL products contained low but detectable CD11c^+^ or CD16^+^ cells (Figure 2E; Figure S1B). Thus, well-expanding TIL products contain high numbers of immune cells, of which the majority consists of T cells.

Interestingly, when we compared the composition of CD3^+^ T cell subsets in the tumor digest *ex vivo* with that of the expanded TIL products, we observed that the distribution between CD8^+^ T cells, Tconv cells and γδ T cells was similar (Figure 1E; Figure 2F; Table S1), indicating that the three T cell subsets expanded in a similar fashion (Figure 2G). We next compared the TIL composition from the neuroblastoma lesions to TILs from adult solid tumors (melanoma, NSCLC, RCC). Even though adult TILs also contained γδ T cell infiltrates *ex vivo*, as previously reported^37–39^, their percentages were lower than in pediatric neuroblastoma lesions (Figure 2H). Furthermore, γδ T cells derived from adult tumor lesions failed to expand (Figure 2H, 2I). Thus, only γδ T cells from pediatric neuroblastoma expanded well with the REP protocol. In sum, TIL products can be generated from neuroblastoma lesions, yet its efficiency depends on the quantity of the starting material. In addition, the generated TIL products display a similar composition of effector T cells measured *ex vivo*.

### Expanded TILs from neuroblastoma display childhood-specific cytokine expression patterns

To test the functionality of TILs that we expanded from neuroblastoma lesions, we focused on the well-expanded TIL products. All CD8^+^ TILs, Tconv TILs, and γδ TILs were potent producers of TNF after 7h stimulation with PMA/Ionomycin (Figure S2A, S2B). However, the production of IFNγ was highly variable for all three T cell subsets (Figure S2A, S2B). Most strikingly, the production of IL-2 upon PMA/Ionomycin stimulation was below 20% for CD8^+^ T cells and γδ T cells, and only four TIL products contained >20% IL-2 producing Tconv cells (Figure S2A, S2B). This limited IL-2 production corroborates with previous reports from blood-derived pediatric T cells^40,41^. Thus, neuroblastoma-derived TIL products can produce cytokines, but do so with lower potency than TILs from adult solid tumors, which show very high production of all three cytokines with PMA/Ionomycin stimulation^18–20^.

### Tumor-reactivity from neuroblastoma TIL products primarily originates from γδ T cells

We next tested whether the well-expanded TIL products contained tumor-reactive T cells. We exposed the TIL products for 6-7h to autologous tumor digest, and we measured the expression of the activation marker CD137 (4-1BB) indicative of TCR triggering^42^, the degranulation marker CD107α, and TNF and IFNγ required for effective anti-tumoral responses^43^ (Figure 3A). To increase the low HLA expression cells of neuroblastoma cells^9^, and thus their capacity to present antigens to T cells, we pre-exposed the tumor digests overnight with 1000 IU/ml human recombinant (hr)IFNγ, prior to co-culture with the TILs.

**Figure 3.**
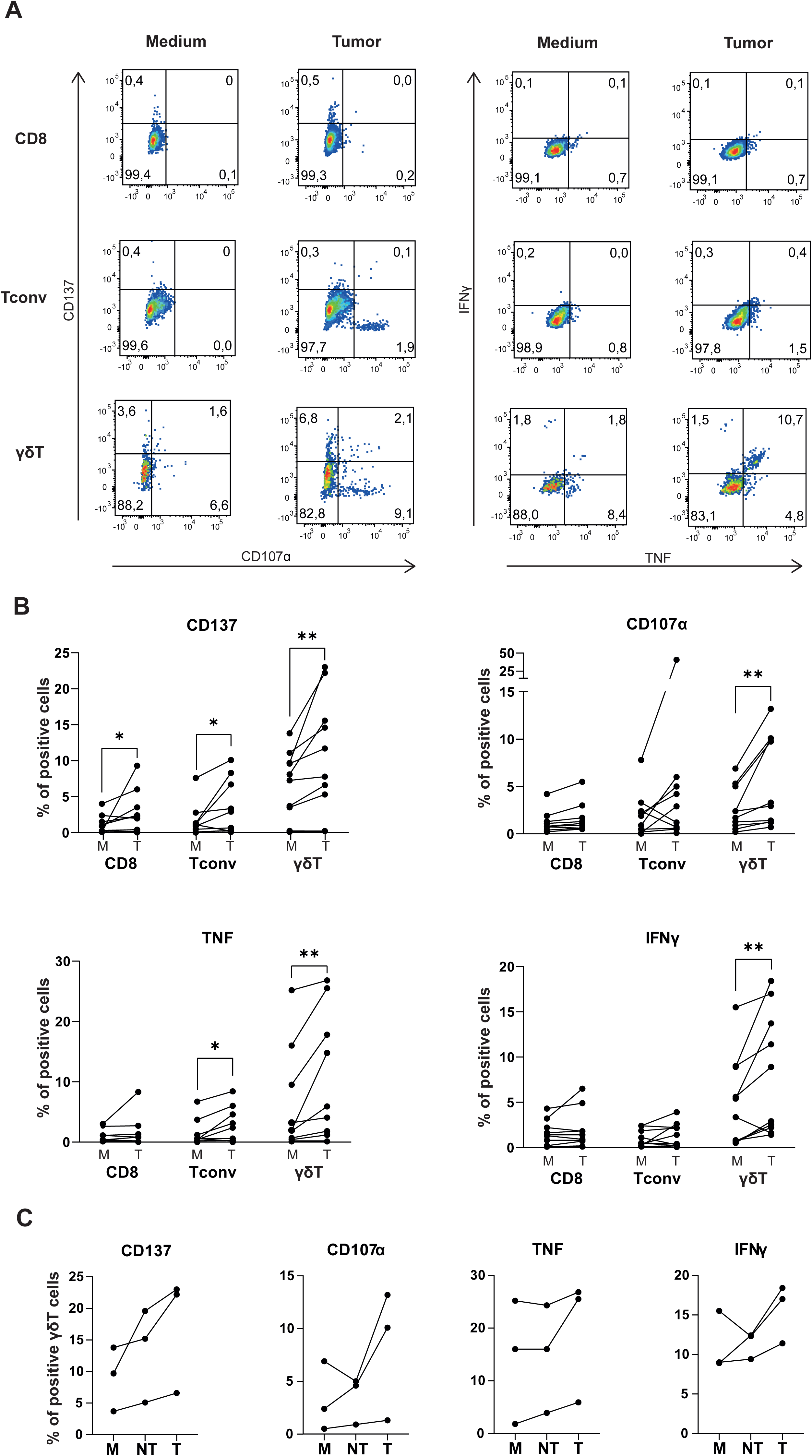
γδ T cells display anti-tumor reactivity against autologous tumor digest. **(A)** Representative dot plots for expression levels of surface molecules (left) and cytokines (right) upon 6-7h culture with medium alone, or with autologous tumor digest in a 1:1 ratio (sample M583AAB). **(B)** Compiled data of CD137, CD107α, TNF and IFNγ expression on indicated T cell subsets after culture with medium (M) or with autologous tumor digest (T) **(C)** Compiled data of CD137, CD107α, TNF and IFNγ expression of γδ TILs after a 6-7 hour co-culture with medium (M), with non-tumorous tissue digest (NT), or with autologous tumor digest in a 1:1 ratio (T). (n=3). Each dot represents one TIL product (n=10). Box and whisker plots depict median, minimum and maximum values (whiskers) and 25th pct and 75th pct (box). Paired student t-test; *<0.05, **<0.01.

Co-culture of TILs with autologous tumor digest resulted in significantly increased CD137 expression in all T cell subsets when compared to medium control (Figure 3B; CD8^+^ T cells: p=0.02; CD4^+^ T cells: p=0.03; γδ T cells: p=0.004). This finding indicates that the TCR was engaged. In contrast, CD8^+^ T cells and Tconv cells displayed very limited expression of CD107α or production of IFNγ, if present at all (Figure 3B). TNF production was also nearly absent in CD8^+^ T cells and limited, yet significantly increased­, in Tconv cells (Figure 3B; p=0.04). In contrast, γδ TILs not only significantly increased CD107α expression when exposed to tumor digest (Figure 3B; p=0.002), but also substantially increased the production of TNF and IFNγ (Figure 3B; TNF: p=0.008; IFNγ p=0.01). Of note, the observed anti-tumor response of γδ TILs did not depend on pretreating tumor digests with hrIFNγ (Figure S2C), suggesting that γδ TILs do not react to Interferon-gamma response genes. Importantly, the observed γδ T cell responses from neuroblastoma-derived TIL products were at least in part tumor-specific, as lower responses were observed to autologous non-tumorous tissue digest cultures than to tumor digest (Figure 3C).

PD-1 and CD137 are two expression markers reported to enrich for tumor-reactive T cells^43,44^. Both markers were expressed on a subset of γδ T cells *ex vivo* (Figure S2D). Whereas PD-1 expression was limited, CD137 expression was significantly higher on γδ T cells than on αβ T cells (Figure S2D). Nevertheless, neither the percentage of PD-1 nor of CD137 *ex vivo* correlated with tumor reactivity of γδ T cells in the TIL product (Figure S2E). When we correlated the response rates of tumor-reactive γδ TILs (e.g. any γδ TIL that upregulated at least one of the four measured markers), we observed a positive trend with the patient’s age, but not with the percentage of γδ T cells *ex vivo* (Figure S2E). Thus, γδ T cells are the most tumor-reactive T cell subset in neuroblastoma lesions, however our patient cohort is too small for identification of markers that could predict tumor-reactivity.

### Vδ1 and Vδ3 γδ T cells are the prime source of anti-tumor responses

We next studied which γδ T cell subset contributed most to the anti-tumor response. γδ T cells can be divided into subclasses based on the Vδ gene used in their TCR^45–47^. In humans, Vδ1, Vδ2 and Vδ3-expressing γδ T cells are the most common subtypes^46,48^. We first measured the γδ T cell composition of four expanded TILs with known anti-tumoral responses (Figure 4A). The Vδ1 subtype was most prevalent in three TIL products, and one primarily contained Vδ2 cells (Figure 4B). The percentage of Vδ3 cells was low but present in all four TIL products (Figure 4B). All γδ T cell subsets were equally equipped to produce TNF and IFNγ when stimulated with PMA/Ionomycin (Figure 4C, 4D). However, when we repeated the co-culture with autologous tumor digests, increased CD137 expression was primarily found on Vδ1 cells and Vδ3 cells, and much less so on Vδ2 cells (Figure 4E, 4F). Increased CD107α expression was found on all γδ T cell subsets but was most prominent on Vδ3 cells (Figure 4F). Cytokine production was also primarily detected on the Vδ1 and Vδ3 cells (Figure 4F). Thus, Vδ1 and Vδ3 cells are the prime γδ T cell subsets responding to autologous tumor digests.

**Figure 4.**
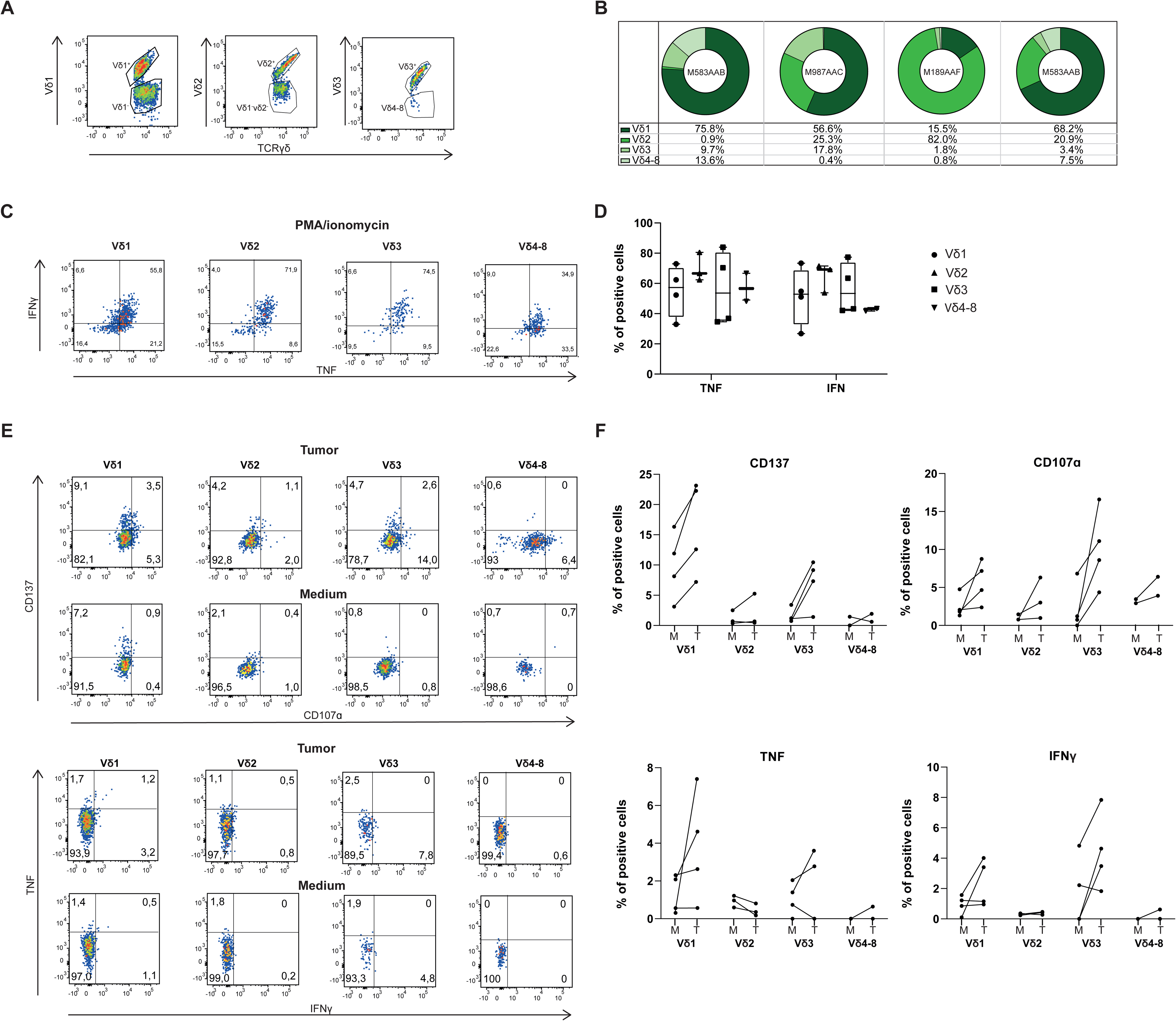
Tumor-reactive γδ T cells primarily consist of the Vδ1 and Vδ3 subset. **(A)** Representative dot plot for Vδ1, Vδ2, Vδ3 and Vδ4-8 TILs, pre-gated on γδ T TILs (sample M987AAC). **(B)** Distribution of Vδ1, Vδ2, Vδ3 and Vδ4-8 as percentage of total γδ T cell population in four expanded TIL products. γδ T cell subsets were excluded from further analysis when cell numbers were <100 cells. **(C)** Representative FACS plots for cytokine secretion by Vδ1, Vδ2, Vδ3 and Vδ4-8 TILs after stimulation for 6-7 hours with PMA-Ionomycin. **(D)** Compiled data (n=4). **(E)** Representative FACS plots and (**F**) of compiled data of indicated marker on Vδ1, Vδ2, Vδ3 and Vδ4-8 TILs after 6-7 hours co-culture with autologous tumor digest in a 1:1 ratio. Each dot represents one TIL product. Box and whisker plots depict median, minimum and maximum values (whiskers) and 25th pct and 75th pct (box).

In summary, we here provide first evidence that TIL products can be generated from pediatric neuroblastoma tumor lesions. Importantly, γδ T cells are the prime anti-tumoral T cells in these TIL products, which show higher response rates to the autologous tumor than non-tumorous tissue digest. It is yet to be determined how γδ T cells recognize neuroblastoma cells. The γδ TCR does not have the obligate HLA restriction as the conventional αβ TCR^49^, and γδ T cells can kill target cells upon engagement of NK-cell receptors^50^. Recent studies reported that γδ T cells respond to checkpoint inhibition in HLA-deficient colon cancer and in melanoma patients with low mutational rate^51,52^. Also, we could not find substantial differences in anti-tumoral responses by γδ T cells to autologous tumors that were pre-treated with hrIFNγ or not, suggesting that the anti-tumoral response of γδ T cells in neuroblastoma is indeed HLA-independent. Of note, evidence that γδ T cells are contributors to anti-tumoral responses in solid tumors is recently further accumulating. Indeed, γδ T cells were shown to be predictive for overall increased survival in non-small cell lung cancer and triple negative breast cancer patients^38,39^.

Importantly, we show that the composition and responsiveness of TIL products from neuroblastoma substantially differs from that of TIL products generated from adult tumors. γδ TILs expand equally well as the αβ T cell subsets, a feature that was not found in three types of adult solid tumors. This could be related to the inherent higher expansion capacity of γδ T cells early after birth^53–55^. In neonates and young children, γδ T cells are more diverse^47^. As children age, γδ T cells become more oligoclonal, decrease in quantity and lose their capacity to expand^56–58^.

Previous studies reported that adult γδ T cells may require other cytokines or stimuli in addition to IL-2 for sufficient expansion^59,60^. Even though we cannot exclude that additional stimuli may even further boost the expansion of γδ T cells, we here show that for pediatric γδ T cells, IL-2 suffices for expansion. Neuroblastoma patients are overall young and mostly below the age of 10, and this may therefore support the effective expansion of γδ T cells with IL-2 alone in our study. In sum, we provide first insights into the difference of adult versus pediatric solid tumor-derived γδ T cells. Based on our findings, we propose that γδ T cells should be considered for treating pediatric solid tumors, as we showcased here for the feasibility to develop autologous, tumor-reactive TIL products.

## MATERIALS AND METHODS

### Patient characteristics

Between February 2020 and January 2022, 19 neuroblastoma patients aged 8-139 months (average: 49.6 months), 11/19 (58%) female, were included in this study. One patient sample (M502AAD) was excluded due to the suspicion of not containing exclusively tumor material but being a lymph node and therefore containing many non-tumor specific immune cells. The included patients’ characteristics, origin of tumor, pre-treatment regimens are reported in Table S1. Patients were categorized as high risk (HR; n=12), medium risk (MR; n=5) and low risk (LR; n=1). Tumors were resected from adrenal gland (n=11), soft tissue (n=4), or other organs (n=3). One patient (M635AAA) was included in the observation group and received no treatment prior to debulking surgery. All other patients had received additional treatment before resection, including standard chemotherapy (n=11), combinations of different chemotherapy regimens (n=5), combination of chemotherapy with MIBG (n=2), or chemotherapy followed by autologous stem cell transplantation (n=1). The study was conducted according to the guidelines of the Declaration of Helsinki and approved by the Institutional Review Board (or Ethics Committee) of the Princess Maxima Centrum for Pediatric Oncology (Utrecht, the Netherlands) (protocol code MEC-2016-739, and date of approval 13 December 2016). Informed consent was obtained from all subjects involved in the study.

### Sample collection

Tumor tissue was obtained directly after surgery and processed within 4 hours. Tumor samples (n=20; from patient M909AAA and M189AAF, 2 samples from different tumor regions were included) were minced into pieces of 1 mm^3^ and digested with collagenase IV (Worthington) for max 1 hour at 37°C and filtered over a 70µm cell strainer to obtain a single cell suspension. Tumor digest was frozen in liquid nitrogen in 90% FCS/10% Dimethyl Sulfoxide (DMSO) (Corning) until further use.

### TIL expansion

Tumor samples were thawed in 50 ml warm RPMI medium (Gibco) supplemented with 2% fetal calf serum (FCS) (Bodego, Bodinco BV), washed, and cultured in 20/80 T-cell mixed media (Miltenyi) containing 5% FCS, 5% human serum (HS) (Sanquin), 1.25 mg/ml fungizone and 6000 IU/ml IL-2 (Proleukin, Novartis). Depending on cell density and cloudiness, cells were cultured in a 24-well (1 ml), 48-well (500 µl) or 96-well plate (200 µl) at 37°C and 5% CO_2_ for 12-14 days, with approximately at a concentration of 1 million cells per ml. Medium was usually refreshed on day 6, 9 and 12, and cells were split when a monolayer of cells was visible in the entire well. After 12-14 days, cells were expanded using a down-scaled minor adjusted version of the clinically approved rapid expansion protocol^15^. Briefly, cells were collected and manually counted (hemocytometer) with trypan blue solution (Sigma). 300.000 viable cells (100.000/well) were cocultured with 5x10^6^-10x10^6^ irradiated peripheral blood mononuclear cells (PBMCs) pooled from 15 healthy donors (feeder cells) in a 24-well plate, containing 30 ng/ml anti-CD3 antibody (OKT-3, Miltenyi Biotec) and 3000 IU/ml hrIL-2 for 12-14 days at 37 C and 5% CO_2_. Medium was refreshed every other day, and cells were split when a monolayer in the entire well was achieved. After 12-14 days, cells were harvested and manually counted. Expanded cells were cryo-preserved in RPMI-1640 medium containing 10% DMSO, 40% FCS until further use.

### Tumor reactivity assay

From the well-expanded TIL products, cryopreserved tumor digest and REP TILs were thawed and incubated in 20/80 T-cell mixed media containing 5% FCS, 5% HS, 1.25 mg/ml fungizone and 500 IU/ml IL-2 overnight at 37°C to recover from thawing. Additionally, tumor digest was stimulated with 100-1000 U/ml IFNγ (bio-techne, R&D Systems) overnight, to increase HLA expression. The next day, cells were counted manually with trypan blue solution. A total of 1x10^5^ live expanded TILs were co-cultured with 1-2x10^5^ live tumor digest cells for 7 hours at 37°C. As controls, 1x10^5^ live expanded TILs were stimulated with 10 ng/ml phorbolmyristate acetate (PMA) (Sigma-aldrich) and 1 µg/ml ionomycin (Sigma-Aldrich) or were cultured with T-cell mixed media only. After 1 hour of co-culture, 1x Brefeldin A (Invitrogen) and 1x Monensin (Invitrogen) and anti-CD107α BUV395 (BD Biosciences) were added.

From three patient tumor digest samples, adherent cell cultures could be successfully grown, however, these tissue cell cultures did not contain tumor cells based on single nucleotide polymorphism (SNP) data. Expanded TILs were co-cultured with single cell non-tumorous tissue cells pretreated overnight with hrIFNγ overnight.

### Flow cytometry analysis

Flow cytometry analysis was performed on defrosted and washed material. Tumor digest ex-vivo was stained with Fixable Viability Dye eFluor™ 506 (ThermoFisher, 65-0866-14) for 20 min at 4°C. After washing, cells were stained with the fluorescently labelled surface antibodies (Table S2) in the dark for 20 min at 4°C. After surface staining, cells were fixed and permeabilized using the FOXP3/Transcription factor staining buffer set (Invitrogen) and subsequently stained with intracellular antibodies for 30 min at 4°C in the dark. Defrosted peripheral blood mononuclear cells (PBMCs) from thirteen adult healthy donors were used for staining controls. Samples were acquired on an Aurora flow cytometer (Cytek Biosciences) with SpectroFlo Software.

Flow cytometry analysis of expanded TIL products was performed with defrosted material. Cells were stained with antibodies against CD3, CD4, CD8, CD56, TCRγδPAN, and additionally with Vδ1, Vδ2, and Vδ3 for analysis of γδ T cells subsets in a separate experiment. For phenotypic analysis, antibodies against CD11c, CD16, CD19, CD20, CD25 and CD45 were added as well for 30 minutes at 4°C in the dark (Table S2). Near-IR was added for dead cell exclusion. Cells were washed and fixed with Perm/Fix Foxp3 staining kit (Invitrogen) according to manufacturer’s protocol. Cells were stained with antibodies against TNF, IFN, CD137 and IL2 (T cell activation) or Foxp3 (phenotypic analysis) for 30 minutes at 4°C in the dark. Cells were washed twice and passed through a 70-µM single-cell filter prior to acquisition with the Symphony A5 flow cytometer (BD Biosciences) or the ID7000 Spectral Cell Analyzer (Sony Biotechnologies). A standardized cryopreserved PBMC sample pooled from four healthy donors was included as control to each measurement. Flow cytometry settings were defined for each patient with single antibody staining. Cell subsets were excluded for further analysis when the cell numbers were below 100 cells, e.g. when their percentage was below 1% of the parent population. Data analysis was carried out with FlowJo Star 10.7.1 (BD, Ashland, OR).

### Statistical analysis

Statistical analysis was carried out with GraphPad Prism 8.0.2 (Dotmatics, San Diego, CA). Data are shown as single data points with box and whiskers showing maximum, 75th percentile, median, 25th percentile and minimum, or as paired data points for each patient sample. Significance was calculated with paired student t-test with a two-tailed P value, variance was calculated as standard deviation. The P value cut-offs were set on *<0.05, **<0.01, ***<0.001, ****<0.0001. If not significant, P value is not shown. Correlation plots were corrected for multiple comparison (n=6) using the Bonferroni method.

## ACKNOWLEDGEMENTS

We like to thank the flowcytometry facility staff and the department of cryobiology from Sanquin, and the nursing staff and other employees involved from the Princess Máxima Center. We thank Dr. John Haanen, Dr. Koen Hartemink, Dr. Kim Monkhorst and Dr. Axel Bex from the Antonius van Leeuwenhoek ziekenhuis-Netherlands Cancer Institute for supporting our studies in adult solid tumors, and Aurélie Guislain for technical support. We also thank all patients and their parents for their willingness to contribute to science. This study was supported by an internal grant of Sanquin (PPOC 21-07) to MCW, by Oncode Institute to MCW, and by Stichting Kinder Kankervrij (KIKA 404 and KIKA 491) to JW and MCW. The authors have no conflicting financial interests.

