## Supplemental figure 1 for "γδ T cells are the prime anti-tumoral T cells in pediatric neuroblastoma"

**A**

| Population | Marker | % of | Mean | Min. | Max. |
|---|---|---|---|---|---|
| Live cells | Trypan blue$^-$ | all cells | 32.7 | 6.0 | 72.0 |
| Immune cells | CD45$^+$ | Live cells | 31.4 | 0.9 | 79.2 |
| T cells | CD3$^+$ | CD45$^+$ | 62.0 | 19.5 | 82.6 |
| Monocytes | CD14$^+$ | | 10.0 | 0.5 | 40.2 |
| B cells | CD19$^+$ | | 4.9 | 0.0 | 37.2 |
| NKT cells | CD3$^+$CD16$^+$CD56$^+$ | | 3.6 | 0.0 | 13.5 |
| NK cells | CD3$^-$CD16$^+$CD56$^+$ | | 5.2 | 0.0 | 19.5 |
| CD8 T cells | CD8$^+$ | CD3$^+$ | 44.0 | 27.3 | 69.2 |
| Tconv | CD4$^+$Foxp3$^-$CD25$^-$ | | 38.2 | 22.3 | 63.6 |
| Tregs | CD4$^+$Foxp3$^+$CD25$^+$CD127$^-$ | | 1.0 | 0.0 | 9.1 |
| γδT cells | γδT PAN$^+$ | | 11.9 | 0.9 | 36.0 |

**B**

| Population | Marker | % of | Mean | Min. | Max. |
|---|---|---|---|---|---|
| Live cells | Trypan blue$^-$ | all cells | 65.8 | 33.0 | 95.0 |
| Immune cells | CD45$^+$ | Live cells | 45.5 | 0.0 | 91.7 |
| T cells | CD3$^+$ | CD45$^+$ | 93.3 | 71.6 | 99.9 |
| Monocytes | CD11c$^+$ | | 1.7 | 0.0 | 10.0 |
| B cells | CD19$^+$ | | 0.2 | 0.0 | 0.7 |
| NK cells | CD3-CD16$^+$ | | 0.7 | 0.0 | 4.7 |
| CD8 T cells | CD8$^+$ | CD3$^+$ | 54.1 | 5.2 | 95.4 |
| Tconv | CD4$^+$Foxp3$^-$CD25$^-$ | | 27.5 | 0.2 | 87.7 |
| Treg | CD4$^+$Foxp3$^+$CD25$^+$CD127$^-$ | | 0.1 | 0.0 | 0.3 |
| γδT cells | γδT PAN$^+$ | | 10.0 | 0.0 | 42.1 |

**C**

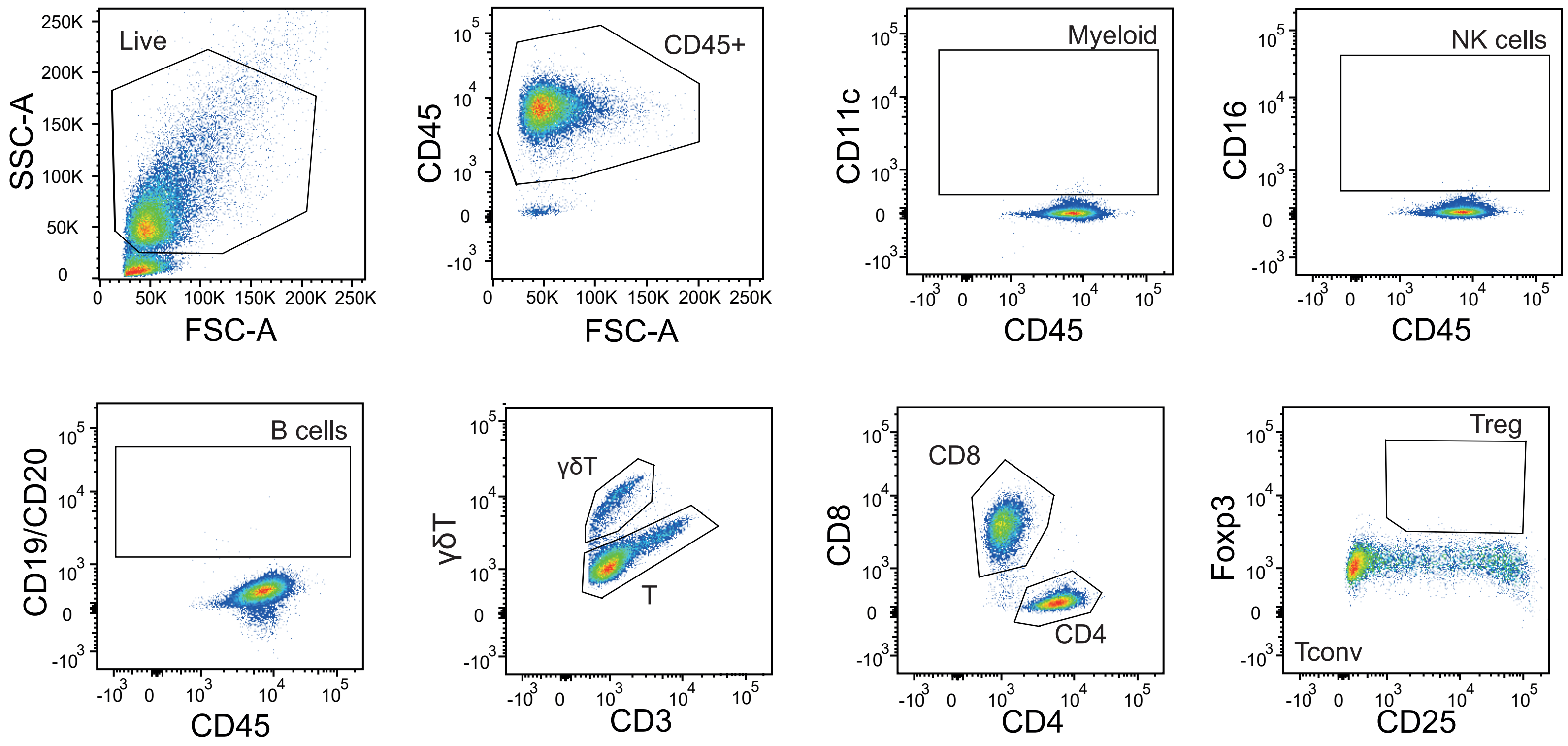


**D**

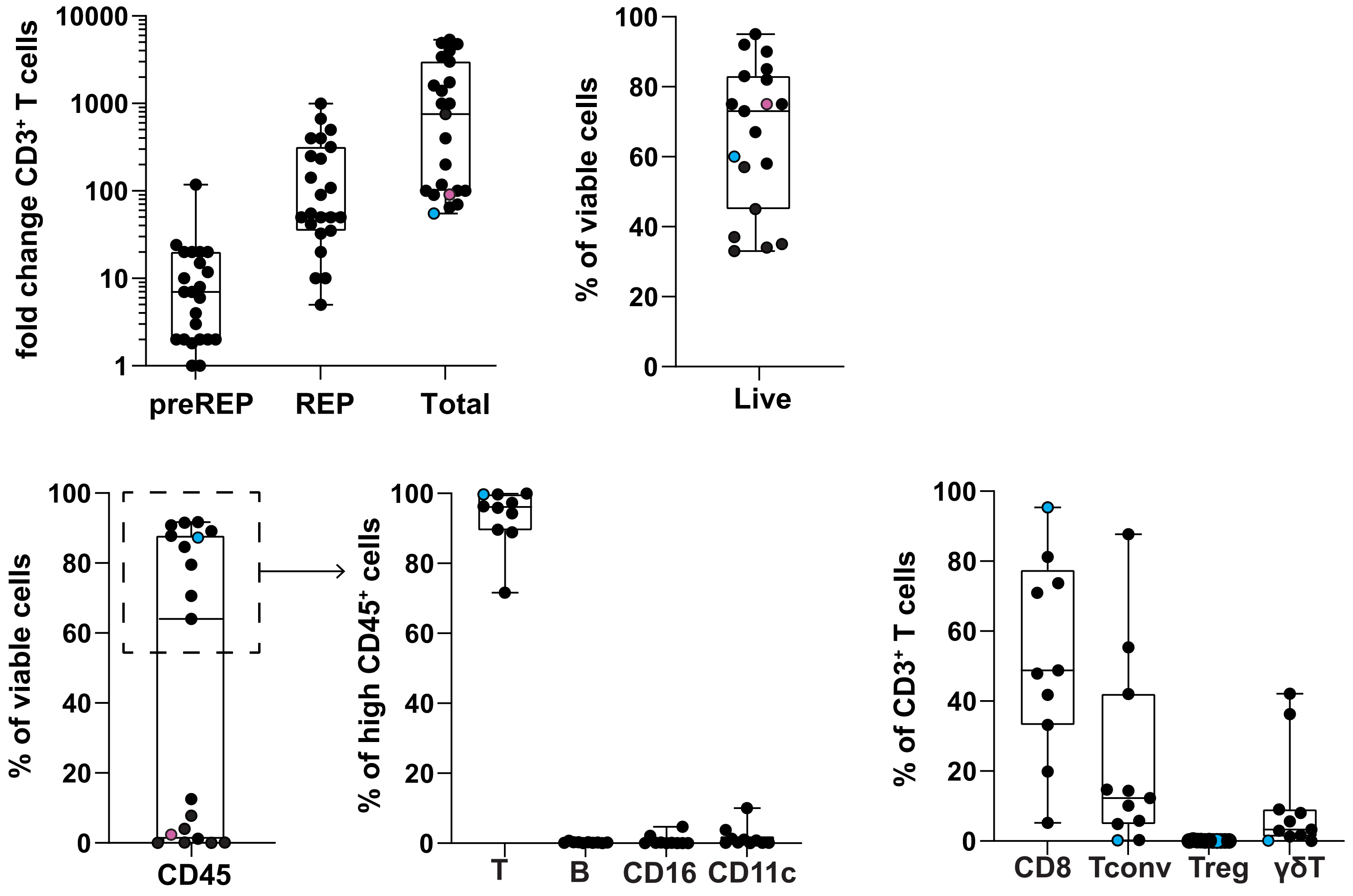
