## Supplementary figures and images for "γδ T cells are the prime anti-tumoral T cells in pediatric neuroblastoma"

### Supplemental figure 2

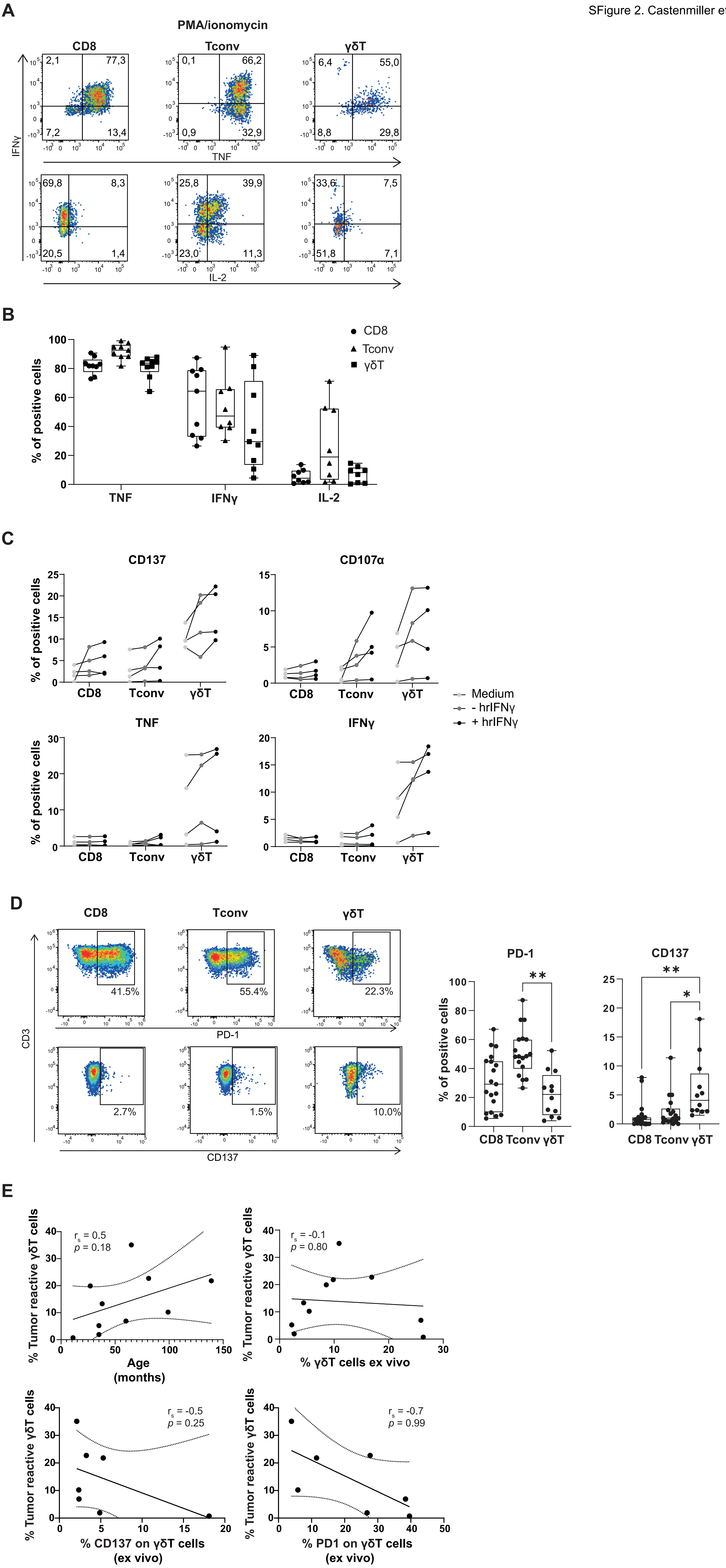
